# MetaPro: A scalable and reproducible data processing and analysis pipeline for metatranscriptomic investigation of microbial communities

**DOI:** 10.1101/2021.02.23.432558

**Authors:** Billy Taj, Mobolaji Adeolu, Xuejian Xiong, Jordan Ang, Nirvana Nursimulu, John Parkinson

## Abstract

**Background:** Whole microbiome RNASeq (metatranscriptomics) has emerged as a powerful technology to functionally interrogate microbial communities. A key challenge is how best to process, analyze and interpret these complex datasets. In a typical application, a single metatranscriptomic dataset may comprise from tens to hundreds of millions of sequence reads. These reads must first be processed and filtered for low quality and potential contaminants, before being annotated with taxonomic and functional labels and subsequently collated to generate global bacterial gene expression profiles.

**Results:** Here we present MetaPro, a flexible, massively scalable metatranscriptomic data analysis pipeline that is cross-platform compatible through its implementation within a Docker framework. MetaPro starts with raw sequence read input (single end or paired end reads) and processes them through a tiered series of filtering, assembly and annotation steps. In addition to yielding a final list of bacterial genes and their relative expression, MetaPro delivers a taxonomic breakdown based on the consensus of complementary prediction algorithms, together with a focused breakdown of enzymes, readily visualized through the Cytoscape network visualization tool. We benchmark the performance of MetaPro against two current state of the art pipelines and demonstrate improved performance and functionality.

**Conclusion:** MetaPro represents an effective integrated solution for the processing and analysis of metatranscriptomic datasets. Its modular architecture allows new algorithms to be deployed as they are developed, ensuring its longevity. To aid user uptake of the pipeline, MetaPro, together with an established tutorial that has been developed for educational purposes is made freely available at https://github.com/ParkinsonLab/MetaPro. The software is freely available under the GNU general public license v3.

## INTRODUCTION

Innovations in culture-independent microbiology, coupled with advances in high-throughput DNA sequencing, have profoundly transformed our understanding of the relationships between microbial communities and their environments^1-3^. In the context of human health, it is increasingly apparent that the composition of the intestinal microbiome has a significant impact on many diseases including type I diabetes, inflammatory bowel disease (IBD), obesity, and rheumatoid arthritis^4-10^. Due to technological and financial constraints, microbiome studies have historically relied on marker gene surveys (e.g.16S rDNA sequences); a technology that focuses on community composition but providing limited insights regarding functional capacity^11,12^. More recently, attention has been turning to the use of whole microbiome DNA and RNA sequencing (metagenomics and metatranscriptomics), which yield more meaningful mechanistic insights through broad analysis of microbiome gene content and gene expression^13-19^. These novel methods of analysis are enabled by Next-Generation Sequencing (NGS) platforms such as Illumina’s HiSeq and NovoSeq platforms, capable of generating the millions of sequence reads required to inform on the thousands of genes encoded and expressed by the microbial communities^20^. A significant challenge is how best to process and interpret these rich datasets that can comprise upwards of hundreds of millions of sequence reads per sample.

While a need for fast and effective tools to automatically process metagenomic and metatranscriptomic pipelines has been identified, few tools are available, particularly for metatranscriptomic data. Web-based analysis platforms such as MG-RAST^21^ and COMAN^22^ have limited support for metatranscriptomic analyses, though the scope and customizability of possible analysis is narrow and the scale of analysis is limited by the availability of remote compute resources. Existing locally hosted metatranscriptomic pipelines such as SAMSA2^23^, IMP^24^, and MetaTrans^25^ offer significantly more options than the web-based analysis pipelines but are insufficiently parallelized, limiting their ability to scale to large (e.g. 100+ GB) datasets. Further, they are relatively inflexible, requiring modifications of experimental protocols or intimate knowledge of computer operating systems to install and execute. The HMP Unified Metabolic Analysis Network (HUMAnN3) is a fast and scalable platform that was primarily designed to analyse metagenomic datasets^26^. Its extension to analyse metatranscriptomic data comes with the expectation that paired (i.e. from the same sample) metagenomic data is available, a potential constraint due to sequencing costs.

Here, we present MetaPro, a flexible, portable, massively scalable end-to-end analysis pipeline for processing metatranscriptomic data. MetaPro is designed specifically to be easy to deploy and use. It is written in Python3 and both the pipeline and associated tools are encapsulated in a Docker image, allowing for single-step installation and deployment on both local computers and scientific computing clusters^27,28^. MetaPro also supports an auto-resume feature for subsequent runs of the same data through the pipeline or if the user wishes to re-run a specific stage of the pipeline. MetaPro is designed with the assumption that new bioinformatics tools will be created that will outperform existing tools currently utilized by MetaPro. Thus, to keep MetaPro relevant, the software architecture enables users to swap, remove, and insert new tools where applicable. To demonstrate improved performance and functionality of MetaPro, we benchmark the speed, resource utilization, and annotation capabilities of MetaPro and two state-of-the-art pipelines, HUMAnN3 and SAMSA2, against three complementary metatranscriptomic datasets. To promote user uptake, both for the application of metatranscriptomics to microbial communities, as well as the tool itself, MetaPro features a tutorial mode that takes the user through each step of the processing pipeline for educational purposes and to encourage adoption beyond the computer specialist.

## RESULTS AND DISCUSSION

### MetaPro – A flexible and scalable metatranscriptomics analysis pipeline

MetaPro was developed as a robust pipeline for the reliable analysis of metatranscriptomic datasets. Key design features include the flexibility to incorporate improved tools as they become available, a scalable architecture to facilitate analysis of hundreds of millions of sequence reads, and ease of use such that end-users are able to install MetaPro as a single software package. To enable these features, the MetaPro pipeline utilizes a modular architecture in which different tools are used at different stages of processing with standard inputs and outputs (**Figure 1**).

**Figure 1:**
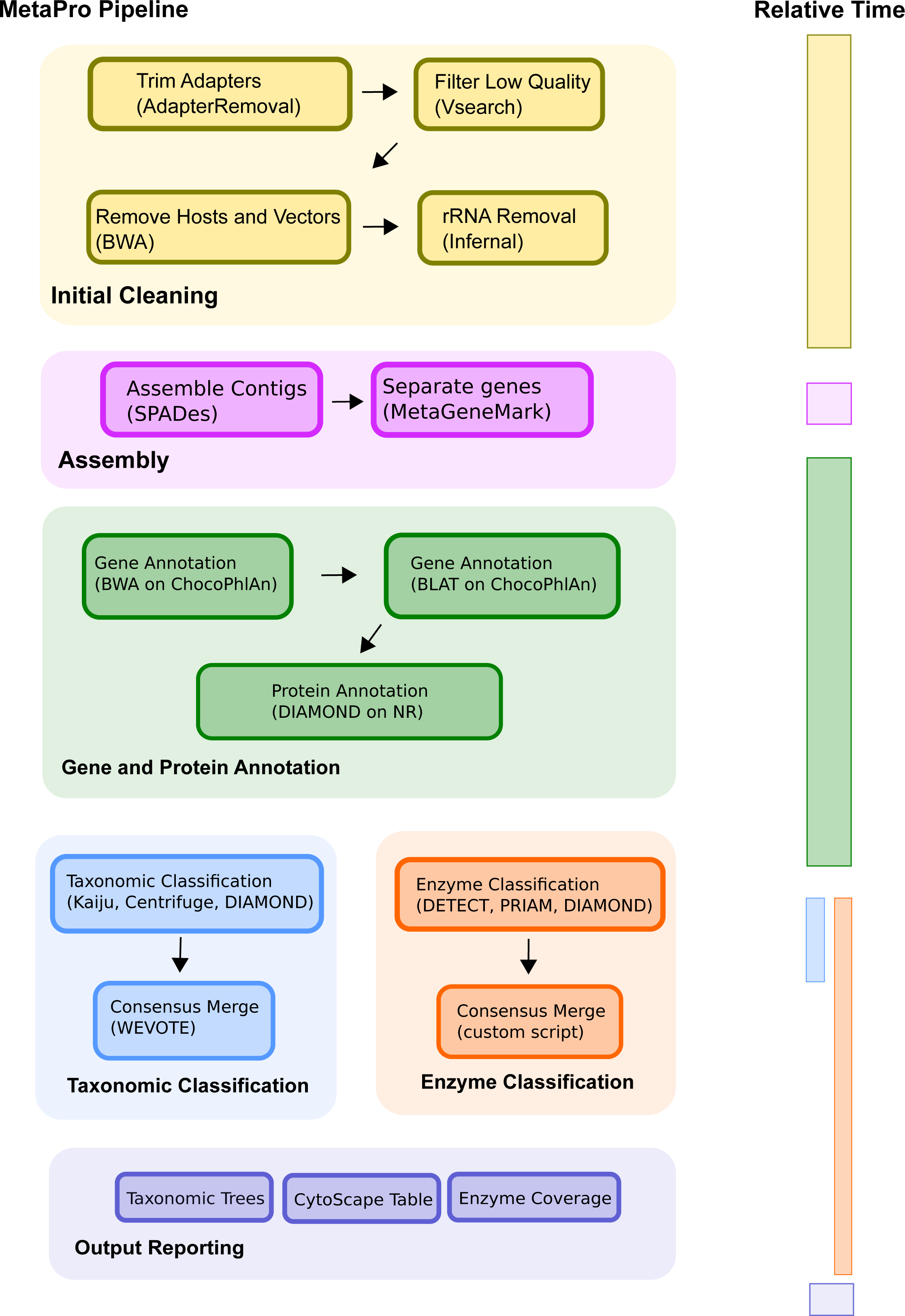
MetaPro Workflow. Overview of MetaPro’s workflow, including the tools and databases used at each step. The bar on the right indicates the relative time required for each phase of the pipeline.

MetaPro accepts demultiplexed FASTQ sequences as the initial input file for analysis. Similar to other pipelines, MetaPro is primarily designed for processing the relatively short-read sequences (e.g. 100-150bp) generated by Illumina platforms, that allow for cost-effective profiling of gene expression in complex microbial communities. MetaPro accepts sequence reads in FASTQ format and first trims and filters for low quality sequences, known host sequences, adaptors, vectors, and both rRNA and tRNA sequences. This results in a set of reads of putative mRNA origin. Previous analyses have demonstrated improved annotation efficiency through assembling reads into longer contigs^29^. Here we apply rnaSPAdes^30^, a transcriptomic assembler within the SPAdes toolkit. Since contigs may feature multiple genes, we subsequently apply MetaGeneMark^31^ to annotate putative genes for each assembled contig. The putative genes and unassembled mRNA reads then undergo three separate multistage processes to annotate: 1) gene identities; 2) taxonomic origin; and 3) enzymatic function.

For gene annotations, MetaPro applies a tiered set of sequence similarity searches, starting with the fastest and least sensitive, BWA^16,32^, followed by pBLAT^33^, then DIAMOND^34^. The former two tools rely on a non-redundant database of genome sequences, ChocoPhlAn^26^, while DIAMOND utilizes the NCBI non-redundant (NR) protein database^35^. For enzyme annotation, MetaPro relies on an ensemble approach involving DETECT^36^, PRIAM^37^ and DIAMOND^34^ searches against the Swiss-Prot database^38^. Due to its greater precision, MetaPro incorporates all DETECT predictions while only incorporating the union of the predictions obtained from both the PRIAM and DIAMOND searches. Finally for taxonomic annotations, MetaPro uses taxonomic assignments of the genes identified through prior searches of ChocoPhlAn and NR protein databases, supplemented with predictions from Kaiju^39^ and Centrifuge^40^, two high-performance short read taxonomic classifiers. These latter classifiers use the NR protein and NCBI nucleotide databases^35^, respectively. Predictions from all sources are combined within a consensus framework to derive a single taxonomic assignment to each contig/read using WeVote^41^

Since we expect MetaPro will be typically deployed on cluster computing environments that feature limitations in memory and/or processing time requirements, MetaPro has been designed to protect against potential points of failure through implementation of intermediate, human-readable, output files that serve as checkpoints. To allow users to readily swap in custom programs and databases to replace currently deployed tools and otherwise extend and modify the pipeline as they see fit, MetaPro is written in Python 3. Further, since ease of installation and use often represent significant barriers to software adoption, MetaPro minimizes software dependencies through the use of the Docker software deployment infrastructure^27^. Docker simplifies the task of distributing pipelines by combining all the tools used by MetaPro in a single image resulting in uncomplicated, single-step installation and deployment independent of computing architecture. Furthermore, MetaPro is fully compatible with Singularity^28^, a secure containerization software based on Docker, that is frequently employed in cluster computing environments.

### Comparisons of MetaPro performance relative to other pipelines

#### Datasets for benchmarking

We benchmarked the performance of MetaPro and two other state-of-the-art metatranscriptomics pipelines, SAMSA2^23^ and HUMAnN3^26^, using three complementary datasets (**Table 1; Supplemental Table 1**). These comprise: 12 samples obtained from the cecum and colon of 5 germ free, non-obese diabetic (NOD) mice inoculated with Altered Schaedler Flora (ASF)^42^ bacteria; 5 samples obtained during a 29 day fermentation of kimchi^43^ at days 7, 13, 18, 25, and 29; and 8 samples obtained during the maturation of an *in vitro* oral biofilm cultured from a complex human oral microbiome^44^ at the 6, 9, 11, 13, 15, 17, 21, and 24 hour time points. The ASF consortium represents a standardized collection of 8 bacteria used in studies of the mouse gut microbiome. The availability of their genome sequence^13^ provides an effective gold standard to benchmark pipeline performance. Similarly, genome sequence data is available for the five species of lactic acid bacteria that, together, dominate over 97% of the kimchi fermentation datasets. Lastly, the human oral microbiome dataset represents putative mRNA reads from a highly complex microbial community encompassing over 700 bacterial taxa, many of which have yet to be cultured. A gold standard is not available for this dataset. However, the intention of analysis of the human oral microbiome dataset was to assess the performance of the three pipelines on a dataset exhibiting similar complexity to microbial communities typically associated, for example, with human health. The performance of each pipeline was assessed against each of these datasets to assess the following metrics: read filtering, gene annotation, taxonomic assignments and enzyme function assignments.

**Table 1:** Sequence read processing efficiency of MetaPro, HUMAnN3, and SAMSA2.

| Dataset | Pipeline | Mean % of total reads that are putative mRNA | Mean % of annotated putative reads | Mean annotated putative reads, as a % of total reads | Mean unique transcripts/gene families* |
| --- | --- | --- | --- | --- | --- |
| NOD mouse | MetaPro | 23.95 | 59.58 | 14.27 | 14377 |
| NOD mouse | HUMAnN3 | 79.95 | 2.80 | 2.24 | 2691 |
| NOD mouse | SAMSA2 | 39.33 | 10.85 | 4.27 | 11499 |
| Kimchi | MetaPro | 32.21 | 59.27 | 19.09 | 87193 |
| Kimchi | HUMAnN3 | 89.26 | 11.71 | 10.45 | 9166 |
| Kimchi | SAMSA2 | 72.54 | 15.72 | 11.4 | 86270 |
| Oral Biofilm | MetaPro | 97.48 | 96.96 | 94.52 | 95707 |
| Oral Biofilm | HUMAnN3 | 100 | 71.31 | 71.31 | 28750 |
| Oral Biofilm | SAMSA2 | 98.98 | 80.40 | 79.58 | 298315 |

\* Unlike MetaPro and SAMSA2, HUMAnN3 reports only the number of unique gene families.

#### Read Filtering

Sequence processing pipelines require the filtering of input read sequences based on the sequence quality and potential contamination. For metatranscriptomics, sources of contamination include sequence adaptors, reads of host origin and RNA reads derived from non-mRNA sources. Identification and removal of these reads is important to both reduce downstream processing time as well as to avoid potentially incorrect inferences of data generated by the pipeline. We therefore examined the performance of each pipeline to filter reads from each of the benchmark datasets (**Table 1**). Filtering of contaminant reads is particularly important in the context of host-associated metatranscriptomic datasets as sequences of host origin can represent a significant proportion of reads in a sample^15^. Both MetaPro and HUMAnN3 (which relies on a Bowtie2-based^45^ sequence filtering tool termed Kneaddata^46^) provide the option to filter host-associated reads. Both pipelines allow for user-defined custom reference databases of host-associated reads to be used in this filtering step. In our performance testing, we utilized the same mouse and human host-associated sequence reference libraries in both tools to ensure comparability between the results. SAMSA2, on the other hand, does not allow for filtering contaminant reads.

Due to the lack of a polyA tail, bacterial RNASeq datasets often feature a high abundance of non-mRNA reads such as those of rRNA or tRNA origin^15^. Identification and removal of these reads can significantly reduce the size of the putative mRNA dataset and increase the speed of downstream analyses. Furthermore, during the generation of many metatranscriptomic datasets rRNA depletion kits are typically employed to reduce these abundant sequence moieties. Such kits often perform better for specific taxa, resulting in taxonomic bias in the removal of rRNA sequences. It is therefore important to bioinformatically filter for any remaining rRNA to minimize their impact on taxonomic readouts. MetaPro filters rRNA and tRNA using a two-tiered approach consisting of the BAsic Rapid Ribosomal RNA Predictor (Barrnap)^47^ and Infernal^48^. Barrnap is a fast RNA filtering tool utilizing hidden Markov models of sequence families, whereas Infernal is a slower, more sensitive RNA filtering tool utilizing both RNA sequence and secondary structure to identify sequence-divergent RNA homologs that conserve their secondary structure. Combining these tools results in sensitive filtering of rRNA and tRNA at an increased speed compared to using only Infernal. In contrast, HUMAnN3 uses the Bowtie2- based KneadData^46^ for rRNA sequence filtering and SAMSA2 uses SortMeRNA^49^, a fast sequence-based filtering tool, to filter rRNA sequences.

Due to the methodological differences discussed above, the three pipelines analysed here each generated differing numbers of putative mRNA reads (**Supplemental Table 1**). When analyzing the NOD mouse dataset, MetaPro, SAMSA2, and HUMAnN3 predicted that an average of 24.0 ± 15.6%, 39.3 ± 11.5%, and 80.0 ± 8.3%, respectively, of the total reads in the sample as being of putative mRNA origin. The significantly higher proportion of predicted mRNA reads in these samples by HUMAnN3 likely represent insufficient filtering of low-quality or host-associated reads. When analyzing the kimchi fermentation dataset, MetaPro predicted that an average of 32.2 ± 12% of the total reads in the sample were putative mRNA while SAMSA2 and HUMAnN3 predicted 72.5 ± 3.9% and 89.3 ± 1.2%, respectively. Like the NOD mouse dataset, the high proportion of predicted mRNA reads by SAMSA2 and HUMAnN3 likely represent instances of insufficient filtering. Lastly, when analyzing the human oral biofilm dataset, which had low quality and putative rRNA reads prefiltered^44^, all three pipelines predicted that nearly all of the sample consisted of putative mRNA reads. MetaPro, SAMSA2, and HUMAnN3 predicted 97.5 ± 1.1%, 99.0 ± 0.4%, and 100.0 ± 0.01% putative mRNA, respectively.

These differences in annotation counts reflects the way in which each pipeline handles their data. SAMSA2 separates the unmerged reads but only annotates the merged reads. This results in far fewer putative reads when compared to MetaPro or HUMAnN3. HUMAnN3 does not merge its reads, but when presented with paired-ended data, treats the forward, and reverse reads as two concatenated sets. Thus, HUMAnN3 has the most putative reads of the pipelines. MetaPro attempts to merge pairs of reads, and subsequently attempts to assemble merged pairs, as well as unmerged singletons into contigs. Contigs, merged pairs and unmerged singletons are subsequently subject to annotation. For read accounting purposes, MetaPro considers the two reads associated with a pair as a single sequence (i.e. both reads are assumed to have originated from the same transcript). Note, for unmerged pairs of reads, this results in each read undergoing annotation independently, but with the pair ultimately receiving only a single annotation.

#### Assigning Reads to Genes

The ability to assign a short read to a gene is a critical step towards functional and taxonomic profiling of metagenomic and metatranscriptomic datasets. Here we assessed the ability of each pipeline to assign reads of putative mRNA origin to individual transcripts. For the NOD mouse and Kimchi datasets, we were interested in comparing each tools ability to map reads to genes associated with the taxa known to be present in these datasets. While MetaPro and SAMSA2 assign reads to distinct genes and proteins, HUMAnN3 reports only on the basis of gene families as defined by UniRef90^50^. This is reflected in the reports of unique transcripts identified in the datasets (**Table 1 and Supplemental Table 1**), where MetaPro and SAMSA2 report similar results, with the exception of the oral biofilm datasets where the larger number of unique transcripts reported by SAMSA2 is likely associated with less stringent criteria used in sequence similarity searches. For HUMAnN3, gene family abundances are split by taxa and reported in terms of reads per kilobase (RPK), which combines matches to members of a gene family normalized for the length of each member matched as well as for sequences that match to multiple reference genes. Thus, to compare to the gene-centric reporting of MetaPro and SAMSA2, we used intermediate files generated by the Bowtie2 and DIAMOND searches in the HUMAnN3 pipeline to map reads to individual genes and proteins. Reads mapping to genes from multiple taxa were allocated equally to each taxon (with the relative contribution of that read assigned to each taxon being a proportion of the number of taxa the read mapped to). Reads that were not subsequently aligned (and reported) to a UniRef90 gene family were removed.

From each pipelines mappings, we compared the number of putative mRNA reads annotated by each pipeline to genes and/or proteins associated with the 8 ASF (NOD mouse datasets) and 5 lactic acid bacteria (kimchi datasets) known to be present in each sample (**Figure 2**). To establish a gold standard for each sample, we used BWA^32^ to perform sequence similarity searches of reads from each sample against a database comprising only the genomes of either the 8 ASF bacteria or the 5 lactic acid bacteria. While we note that sequence data for all 8 ASF and 5 lactic acid bacteria were present in databases used by each pipeline, we expect that the considerably greater taxonomic representation of sequences in these databases would result in false positive taxonomic assignments.

**Figure 2:**
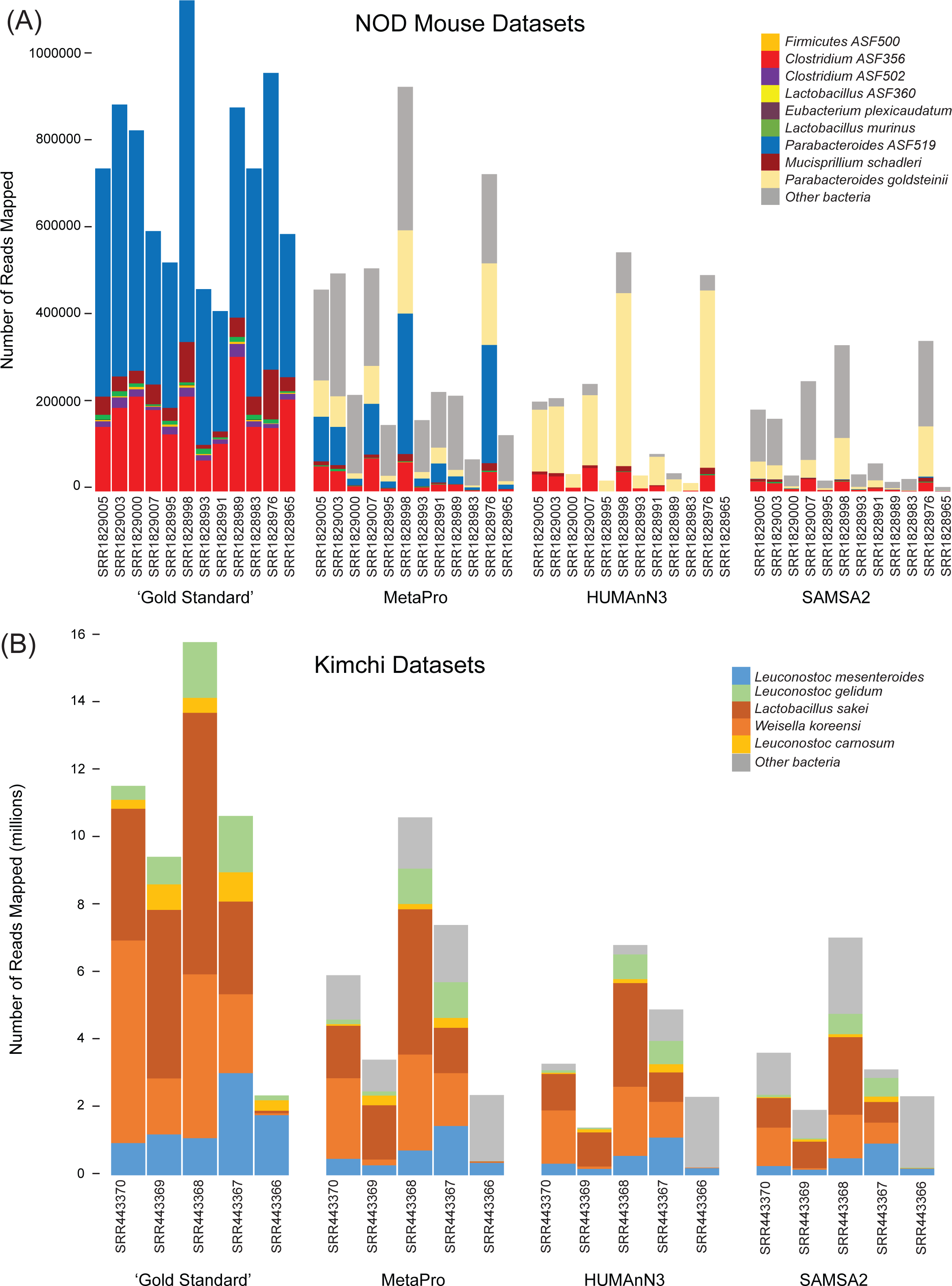
Gene annotation performance of MetaPro, HUMAnN3, and SAMSA2. Stacked barcharts depicting the number of reads annotated to specific taxa in (A) NOD mouse samples, and (B) Kimchi samples by BWA alignments, MetaPro, HUMAnN3, and SAMSA2. The NOD mouse datasets were generated from gut samples from mice inoculated with a defined microbial consortium (Altered Schaedler Flora (ASF); ^42^). In addition to the 8 taxa associated with ASF, reads were also assigned to *Parabacteroides goldsteinii,* a close relative of *Parabacteroides ASF519* (see legend). The kimchi datasets comprise five major taxa (see legend;^75-79^). It should be noted that *Leuconostoc gasicomitatum* reported in the original publication is currently classified as a subspecies of *Leuconostoc gelidum*. For NOD sample SRR1828965, HUMAnN3 did not annotate any reads.

Across the 12 NOD mouse samples, our BWA-based benchmarking annotated a total of 8.6 million reads to genomes of the 8 ASF bacteria (**Figure 2A and Supplemental Table 2**). In contrast to the number of reads annotated to be of putative mRNA origin, MetaPro was able to annotate more reads to a known gene/protein (4.25 million) compared to SAMSA2 (1.49 million) or HUMAnN3 (1.47 million) (**Supplemental Tables 1 and 2)**. Furthermore, MetaPro also annotated a greater number and proportion of these reads to one of the 8 ASF bacteria (1.45 million reads, 34.1%), compared to both SAMSA2 (179,000 reads, 12%) and HUMAnN3 (177,000 reads, 12%). A majority of the reads annotated to ASF bacteria by SAMSA2 and HUMAnN3 were associated with *Clostridium* ASF356. However, *Parabacteroides* ASF519, the most prevalent species in the samples as defined by the gold standard (66.2% ± 6.5% of reads), was absent from the HUMAnN3 results and represented by only 7,053 reads across all 12 samples annotated by SAMSA2. Instead, it appears that both pipelines erroneously assign many reads to transcripts derived from a closely related species, *Parabaceteroides goldsteinii*. In contrast, MetaPro identified 581,849 reads aligning to *Parabacteroides ASF519* in all 12 samples. Furthermore, HUMAnN3 failed to assign any reads for sample SRR1828965 and was unable to assign reads to transcripts associated with *Clostridium* ASF502, *Eubacterium plexicaudatum* ASF492, *Firmicutes* ASF500, and *Lactobacillus* ASF360. MetaPro and SAMSA2 also exhibited poor annotation rates for these taxa, although we note that the version of ChocoPhlAn that MetaPro used in these analyses did not contain *Firmicutes* ASF500.

**Table 2:** Average runtime performance of MetaPro, HUMAnN3, and SAMSA2.

| Dataset | Pipeline | Pre-Processing | St. Dev. | Data Analysis | St. Dev. | Total Time | St. Dev. |
| --- | --- | --- | --- | --- | --- | --- | --- |
| NOD mouse | HUMAnN3 | 339.50 | 101.00 | 970.92 | 211.83 | 1310.42 | 267.13 |
| NOD mouse | SAMSA2 | --- | --- | --- | --- | 2341.75 | 320.92 |
| NOD mouse | MetaPro | 3869.12 | 3442.85 | 12202.28 | 4696.80 | 16071.40 | 5259.01 |
| Kimchi | HUMAnN3 | 1960.80 | 133.92 | 9106.20 | 809.62 | 11067.00 | 902.83 |
| Kimchi | SAMSA2 | --- | --- | --- | --- | 5882.60 | 1738.68 |
| Kimchi | MetaPro | 24853.02 | 3256.61 | 101984.66 | 67339.49 | 126837.68 | 70477.12 |
| Oral Biofilm | HUMAnN3 | 3481.00 | 203.84 | 10800.63 | 967.63 | 14281.63 | 1126.46 |
| Oral Biofilm | SAMSA2 | --- | --- | --- | --- | 53938.88 | 3190.63 |
| Oral Biofilm | MetaPro | 19183.11 | 2479.90 | 41996.21 | 3424.56 | 61179.33 | 5191.15 |

Across the 5 kimchi fermentation samples, BWA-based benchmarking annotated a total of 49.6 million reads to genomes of the 5 lactic acid bacteria (**Figure 2B and Supplemental Table 3**). Similar to the NOD mouse samples, MetaPro was able to annotate more reads to a known gene/protein (19.6 million) compared to SAMSA2 (20.2 million) or HUMAnN3 (18.5 million) (**Supplemental Tables 1 and 3)**. Furthermore, MetaPro also annotated more of these reads to one of the 5 lactic acid bacteria (22.2 million reads, 75%), compared to both SAMSA2 (11.3 million reads, 56%) and HUMAnN3 (15.2 million reads, 82%). Interestingly, sample SRR443366 proved challenging to identify 3 of the 5 lactic acid bacteria: *Weissella koreensis*, *Leuconostic carnosum*, and *Leuconostic gelidum*, relative to other samples. Few reads were annotated to these taxa by both MetaPro and SAMSA2, and no reads were annotated to these taxa by HUMAnN3.

From these analyses, we found that MetaPro outperforms SAMSA2 and HUMAnN3 in terms of assigning putative mRNA reads to the appropriate taxon of origin. For HUMAnN3, the limited ability to assign reads to appropriate taxa may reflect the pipelines reliance on paired metagenomic data and the use of MetaPhlAn3^51^ to identify taxa likely associated with the dataset being processed. This allows the use of smaller, more targeted, reference databases for sequence similarity searches, greatly reducing runtime. However, this strategy may be compromised if the taxa have not previously been well sampled, potentially explaining the inability of HUMAnN3 to annotate any reads for sample SRR1828965.

We note that HUMAnN3 was recently developed to supersede a previous version, HUMAnN2. Comparisons reveal these two versions generally yield similar results (**Supplemental Figure 1; Supplemental Tables 1-3**). However, for the NOD mouse datasets, HUMAnN3 assigned more reads to *Clostridium ASF356* over its predecessor, while HUMAnN2, unlike HUMAnN3, was able to assign at least some reads in sample SRR1828965. For the kimchi datasets, HUMAnN3 was able to assign more reads to *Leuconostoc gelidum*, and *Leuconostoc carnosum* and, unlike HUMAnN2, was able to assign reads in sample SRR443366. We suggest these differences in the ability to assign reads to the two datasets, SRR1828965 and SRR443366 may be related to the different versions of MetaPhlAn used be each pipeline to generate targeted reference databases.

Overall, we found that MetaPro best reflected the gold standard assignments, although as for the other pipelines, MetaPro assigned many reads to sequences from other taxa not expected in the samples. While this highlights the need to improve gene annotation methods, we nonetheless note that such sequences represent potential homologs of genes encoded by the expected taxa and are thus likely to be functionally informative.

#### Inferring Accurate Taxonomic Ranks

In the previous section we benchmarked each pipelines ability to map putative mRNA reads to transcripts of genes encoded by the genomes of bacterial taxa known to comprise the datasets. However, determining the taxonomic composition of a sample to elucidate taxa responsible for providing critical functions remains a challenge in metatranscriptomic analysis. MetaPro features an additional stage for taxonomic assignment to more accurately assign taxonomic classification to putative mRNA reads. MetaPro utilizes input from the short-read classifiers Kaiju and Centrifuge^40^, together with the inferred taxonomy from gene annotations derived from the initial tiered set of sequence similarity searchers, to determine the taxonomic composition of a sample. The taxonomic output from the short-read classifiers and the gene annotation stage are combined into a single consensus taxonomic classification using WEVOTE^41^ in order to increase the precision of classification. WEVOTE considers the taxonomic predictions from the two short-read classifiers, as well as the gene annotations, using a simplified variant of the NCBI taxonomy tree structure and selects the highest confidence taxonomic classification based on the output from individual approaches. WEVOTE produces a consensus classification depending on the pattern of taxonomic classification produced by the three predictors. HUMAnN3 utilizes MetaPhlAn3^52^ for taxonomic profiling based on clade-specific marker genes in the sample whereas SAMSA2 solely utilizes the curated taxonomic annotations present in the RefSeq database to determine read classification.

To assess the performance of each pipeline, we compared the total number of reads that were correctly annotated to specific taxonomic ranks, from species to phylum, within the 8 ASF and 5 lactic acid bacteria for the NOD mouse and kimchi fermentation datasets (**Figure 3**). For the mouse NOD mouse datasets^15^, MetaPro can annotate an average of 55.7 ± 9.2% of reads across all samples, 49.9 ± 8.5% down to the species level, and 2.7 ± 0.4% to the genus level. The remaining 3.1 ± 1.5% of reads were annotated to the rank of family or higher. SAMSA2 annotated 8.1 ± 6.5% of each sample, with 0.2 ± 8.5% down to the species level and 6.3 ± 5.7% to the genus level, while HUMAnN3 annotated an average of 5.6 ± 6.7% reads to each sample only to the species level. HUMAnN3 defaults to reporting on species-level taxa.

**Figure 3:**
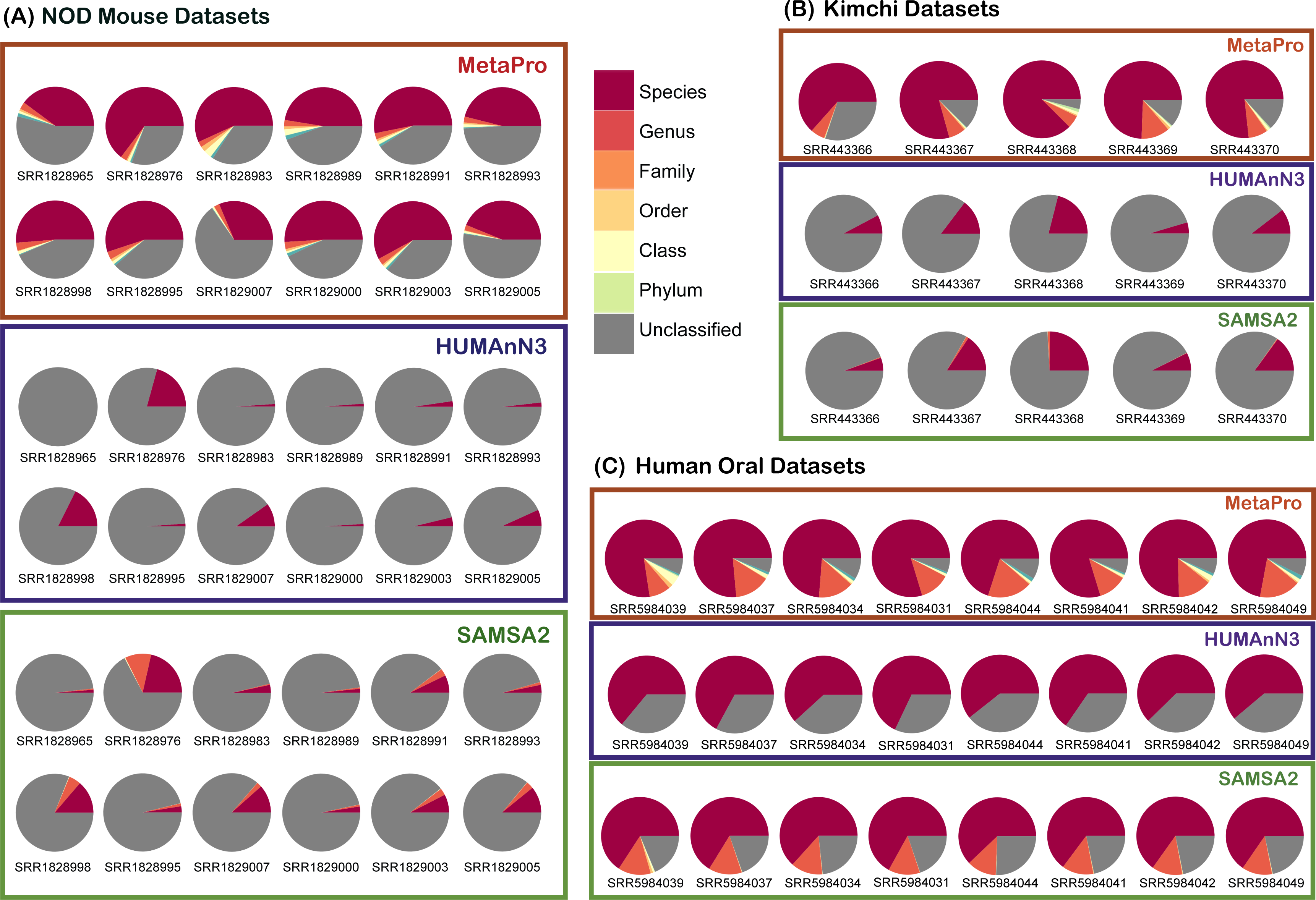
Taxonomic classification performance of MetaPro, HUMAnN3, and SAMSA2. For the NOD mouse (A) and Kimchi (B) datasets, each pie chart shows a breakdown of taxonomic assignments at the different phylogenetic levels indicated, that are closest to the last common ancestor of the expected bacteria within the sample. For the human oral datasets (C), given the lack of gold standard assignments, each pie chart represents the relative abundance of reads assigned to different phylogenetic levels. Unclassified reads represent annotated reads with no assigned taxon.

For the Kimchi set, MetaPro can annotate average of 85.8 ± 8.5% of putative reads across all samples, 76.3 ± 7.8% down to the species level, and 7.6 ± 2.3% down to the genus level. The remaining 1.9 ± 0.9% of the reads annotated to the rank of family or higher. SAMSA2 annotated an average of 13.4 ± 7.6% across all kimchi samples, with 5.2 ± 3.5% to the species level, and 6.59 ± 4.40% to the genus level. The remaining 1.6 ± 0.3% annotate to a taxon of family or higher. HUMAnN3 annotated an average of 11.7 ± 5.7% reads to the species level, and only to the species level.

In addition to the NOD mouse and kimchi fermentation datasets, we were also interested in examining the performance of the pipelines on classifying more complex datasets derived from human oral microbiomes. (**Figure 3C**). These datasets comprise 8 samples and provide more complex data to further assess the performance of the three tools, with the caveat that we do not know what taxa are present in these samples. MetaPro annotated an average of 93% ± 1% of the data, leaving 7% ± 1% unable to be identified SAMSA2 was able to annotate an average of 80% ± 1.9% of its putative reads, with 20% ± 1.9% of the reads unclassified to any known taxa. HUMAnN3 annotated an average of 63.8% ± 2.6% of its putative reads, leaving 36.1 ± 2.6% unmapped. Of MetaPro’s putative reads, an average of 75.6 ± 3.2% were identified down to the species level, and 14% ± 2.8% of the reads were annotated to the genus taxa. SAMSA2 annotated an average of 65% ± 1.5% of its putative reads to a species, and 13% ± 0.5% of those reads to a genus. HUMAnN3 annotated all 64.8 ± 2.6% of its putative reads to a species.

Overall, we found that MetaPro’s ensemble approach to taxonomic annotation consistently outperformed the approaches employed by SAMSA2 and HUMAnN3, although we did note a performance improvement for HUMAnN3 over HUMAnN2 (**Supplemental Figure 2**).

#### Annotating Enzymatic Functions

The ability of each pipeline to accurately infer enzymatic functions of the datasets is essential for understanding the metabolic activity of a sample. Since annotations based on simple similarity searches can yield false positive rates of up to 50%^53^, MetaPro relies on a robust approach that combines predictions from DETECT^36^, with those from PRIAM^54^ that are also confirmed by sequence similarity searches against the Swiss-Prot database^55^. This approach has been shown to significantly outperform sequence similarity searches and has been effectively applied in a number of settings^56-61^

HUMAnN3 reports MetaCyc pathway abundances, but to do this, the gene families are translated from their UniRef IDs into MetaCyc reactions, using a static relational map. This map also includes Enzyme Commission (EC)^62^ numbers. HUMAnN3 then uses a pathway-to- reactions mapping, locating minimally-satisfied paths using MinPath^63^ SAMSA2 uses a static relational mapping of the RefSeq^64^ database for gene annotation, and the SEED subsystem^65^ database for enzymatic function annotation. Both databases are searched using DIAMOND-based sequence similarity searches using a relatively permissive e-value cutoff of 0.001.

MetaPro performs enzyme annotations by first translating genes identified through the BWA and pBLAT searches into proteins and adding these to the proteins identified through the DIAMOND-based searches. Enzyme predictions reported by MetaPro consist of all the DETECT results, together with the intersection of enzymes predicted by both PRIAM and DIAMOND using an e-value cutoff of 1e-5. MetaPro also includes a high-confidence mode that will only return enzymes (reported as ECs) that the pipeline detects with high stringency cutoffs for PRIAM (probability score greater than 0.5) and DIAMOND (e-value cutoff of 1e-10). MetaPro can sometimes predict multiple enzymes per protein. To resolve these instances, MetaPro compares the enzyme matches against a reference database of valid pairs of enzyme combinations compiled from the Swiss-Prot database. Pairs that lack experimental support (i.e. not previously identified in Swiss-Prot) are removed. In situations where more than two enzymes are predicted in a single protein, the enzymes with the two highest probability scores are used.

In all samples, we found that MetaPro identifies more unique ECs than HUMAnN3 and SAMSA2 (**Figure 4; Supplemental Tables 4 & 5**). For example, comparisons of predictions for the NOD mouse samples (**Figure 4A**) reveals MetaPro identifies an average of 949 unique ECs/sample (622 defined as high-quality) compared to 465 and 170 for SAMSA2 and HUMAnN3 respectively. Of these, 350 were shared with SAMSA2 and 153 were shared with HUMAnN3. Both SAMSA2 (41 ECs) and HUMAnN3 (4 ECs) predicted combinations of ECs that have not been identified in the same protein in Swiss-Prot. The kimchi datasets show a similar performance (**Figure 4B**), with MetaPro identifying 1982 unique ECs/sample (1238 defined as high quality) compared to 859 and 600 for SAMSA2 and HUMAnN3 respectively. Of these, 692 were shared with SAMSA2 and 512 were shared with HUMAnN3. For these datasets, SAMSA2 and HUMAnN3 predicted 75 and 32 EC combinations of ECs that have not been identified in the same protein in Swiss-Prot. Finally, for the oral biofilm samples (**Figure 4C**), MetaPro identified an average of 1465 unique ECs/sample (1024 defined as high quality), compared to 858 and 1171 for SAMSA2 and HUMAnN3 respectively. Of these, 674 and 856 were shared with SAMSA2 and HUMAnN3 respectively. Furthermore, SAMSA2 and HUMAnN3 identified an average of 79 and 120 EC combinations not previously seen in Swiss- Prot annotated proteins.

**Figure 4:**
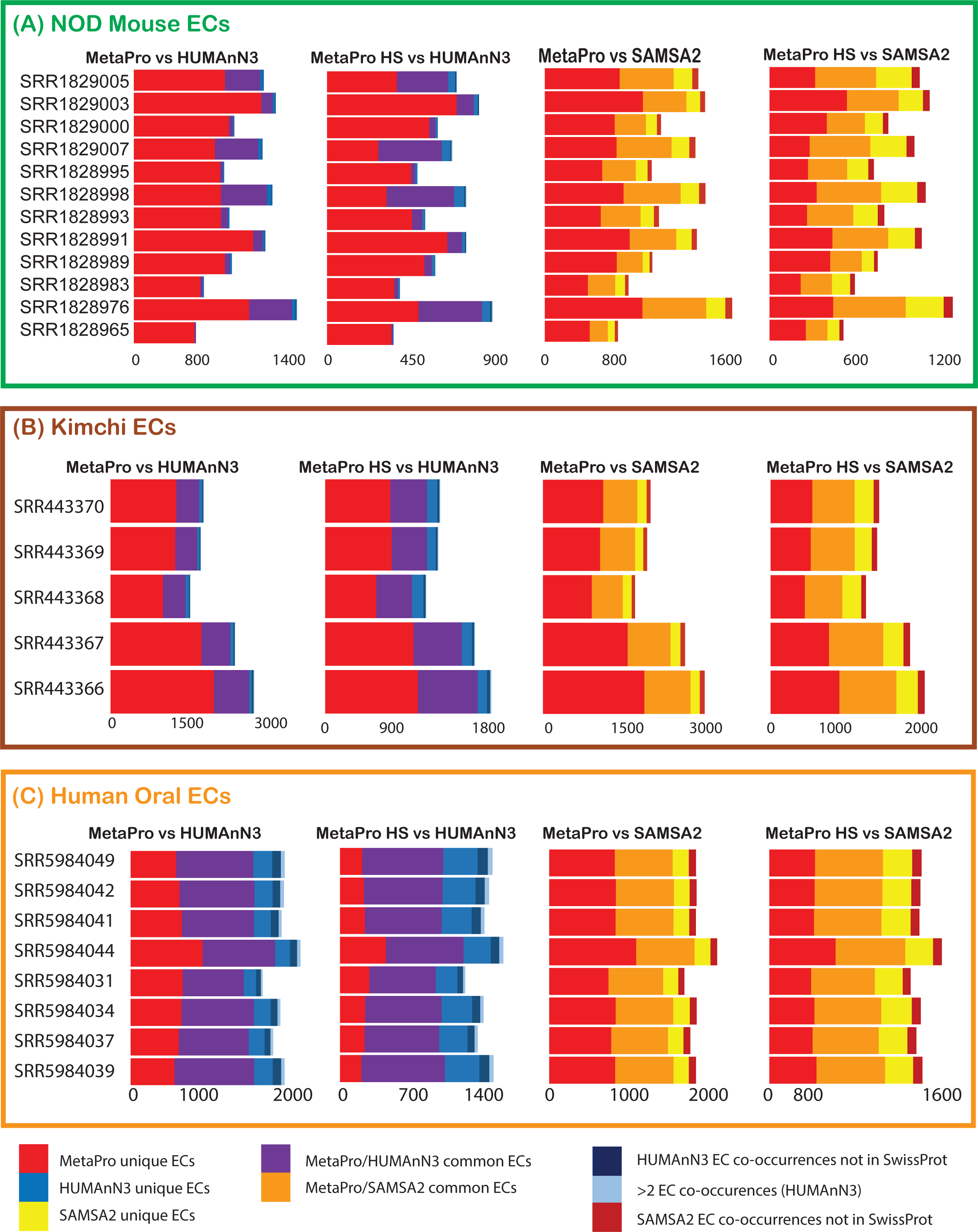
Enzyme annotation performance of MetaPro and HUMAnN3, and SAMSA2. Stacked barcharts indicate the number of enzymes, as defined through enzyme classification (EC) assignments, annotated by each pipeline for the three sets of datasets: (A) NOD Mouse, (B) Kimchi, and (C) Human Oral Biofilm data. In addition to displaying ECs unique or shared between MetaPro and the other two tools, also shown are ECs, predicted by HUMAnN3 and SAMSA2 to occur in combination with another EC, in the same transcript, with no supporting evidence that such a combination has been previously observed (as defined through Swiss-Prot annotations). Further, for HUMAnN3, we show the number of EC assignments that occur in combinations of greater than 2 ECs.

In summary, MetaPro delivers superior performance over the other two tools with both a greater number of EC predictions and greater confidence of annotations (through the combined use of DETECT, PRIAM and DIAMOND predictions) relative to the simple sequence-similarity-based approaches used by the other two tools. As a sidenote, we did find that HUMAnN3 showed improved performance over HUMAnN2, with the latter identifying fewer ECs in both the kimchi and oral biofilm datasets (588 and 986 unique ECs respectively; while also predicting a higher number of combinations of ECs not supported by Swiss-Prot annotations (**Supplemental Figure 3 and Supplemental Table 6**).

### Result Reporting

To allow for more intuitive exploration of processed data, MetaPro features several text and visual outputs. For context, HUMAnN3 produces three text files: 1) a gene families file that details the abundance of each gene family in the community (as measured by reads per kilobase; RPK), stratified to show the contribution from each species; 2) a pathway abundance file that details a normalized abundance of pathway components for each pathway, again stratified to show species contributions; and 3) a pathway coverage table, which provides a confidence score for the presence of a pathway in the community, as well as in individual species. SAMSA2 is typically used to compare between two datasets but does generate files that report: 1) DIAMOND search results; 2) a taxonomic summary of read counts; and 3) an enzyme classification summary of read counts. In contrast, MetaPro produces the following: 1) a histogram of read quality, together with a summary of read processing (reads filtered for quality, host contamination, rRNAs and tRNAs; putative mRNAs; annotated mRNAs; number of unique transcripts; and number of unique ECs); 2) a detailed gene-to-read mapping of every read the pipeline annotated; 3) a summary of every taxon identified in the sample, together with read counts; 4) a list of ECs identified, together with the gene or protein providing the annotation; 5) a table of genes with associated reads per kilobase of transcript, per million mapped reads (RPKM), with details on the contribution of taxa that with further details on the contribution of taxa that are individually responsible for at least 1% of putative reads; 6) a heatmap in .png image format, showing the relative contribution of the 20 most prevalent taxa to enzyme expression grouped in Kyoto Encylcopedia of Genes and Genomes (KEGG)-defined superpathways (**Figure 5A**); and 7) a CytoScape-compatible annotation table that can be readily imported into Cytoscape and mapped on to KEGG defined pathways to illustrate the contribution of specific taxa to enzymes expressed in the selected pathway (**Figure 5B**). This latter mapping requires the use of the KEGGmapper and enhancedGraphics app plugins for the Cytoscape platform and is described in more detail in the accompanying tutorial.

**Figure 5:**
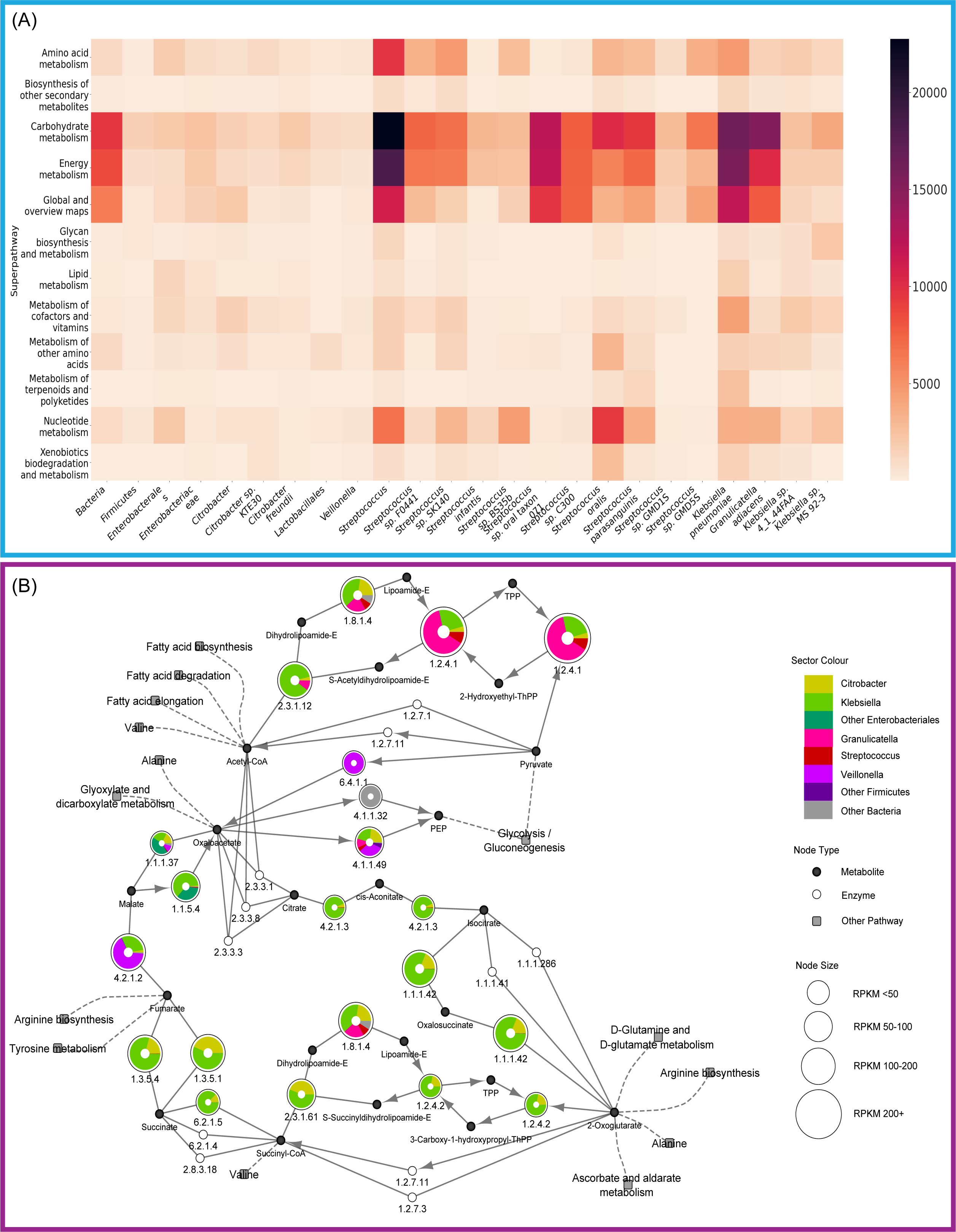
Examples of visual output generated by the MeatPro pipeline. (A) Summary overview showing the contribution of major taxa to KEGG-defined superpathways. After annotation of enzymes, MetaPro generates a heatmap in PNG format showing different taxa responsible for expression (calculated as the sum of RPKMs assigned to each EC annotated for each taxon). Only taxa associated with at least 1% of total reads are shown (see Methods). (B) Cytoscape network-based representation of the tricarboxylic acid (TCA) cycle, together with the breakdown of EC expression by taxon. MetaPro generates a Cytoscape compatible annotation file which can be used to map RPKM data onto KEGG defined pathways using the KEGGscape and enhancedGraphics apps. Here each node represents an individual enzyme with its size indicating the overall expression of that enzyme in the dataset (as defined by RPKM). Coloured sectors indicate the contribution of each taxon to expression of that enzyme. To simplify the display, taxa were manually merged into 8 taxonomic groupings. Here we see that members of Klebsiella and Citrobacter are the main contributors to the TCA cycle. These outputs were generated from a sample in the human oral biofilm dataset (SRR5984039).

### Computational Overhead

Processing time represents an inherent tradeoff between the robustness of an analysis and the speed with which it can be completed. MetaPro, SAMSA2, and HUMAnN3 are each designed to perform specific analyses. MetaPro focuses on the annotation of samples of metatranscriptomic data to provide a detailed breakdown of taxonomic contributions. SAMSA2 was designed to perform comparative analyses between two datasets. HUMAnN3 was primarily designed primarily for the analysis of metagenomic data, reporting gene family abundances, but also accepts metatranscriptomic data. We compared the execution of the MetaPro, SAMSA2, and HUMAnN3 pipelines across all 25 datasets using one computing node equipped with 20 Intel Skylake 2-core-CPU, 202GB RAM, running CentOS 7. Each pipeline differs significantly in their completion times (**Table 2 & Supplemental Table 7**). For the NOD mouse datasets, the mean completion time for each sample was 16071 ± 5259, 2342 ± 321 and 1310 ± 267 seconds for MetaPro, SAMSA2 and HUMAnN3 respectively. Similarly, for the five kimchi fermentation datasets, mean completion time per sample was 53733 ± 18287, 5883 ± 1739 and 11067 ± 903 seconds for MetaPro, SAMSA2 and HUMAnN3 respectively. However, for the 8 human oral biofilm datasets, we found the mean completion times were longer for SAMSA2 being 46835 ± 2679, 53939 ± 3191 and 14282 ± 1126 seconds for MetaPro, SAMSA2 and HUMAnN3, respectively. With the exception of the oral biofilm datasets, MetaPro requires a longer execution time than the other pipelines, with, for example roughly 30% of runtime dedicated to the initial cleaning step, which includes rRNA removal (**Figure 1**). However as indicated above, this increase in runtime stems from the preference to prioritize accuracy over speed. The majority of the execution time within MetaPro is accounted for by both the use of a large reference file (in the case of DIAMOND with the NR database), as well as the use of robust tools for enzymatic function inference (**Figure 1**), rather than static mappings used by other tools. For example, in assigning reads to genes, MetaPro uses the entire ChocoPhlAN database, instead of the subset used by HUMAnN3. Furthermore, MetaPro performs 3 times as many annotation phases as SAMSA2. For the oral biofilm dataset, it was interesting to note the longer execution time of SAMSA2. This appears related to the higher number of reported hits to the RefSeq database during read annotation. This likely resulted in a slow down due to the creation of the large output files associated with reporting the results of the searches.

Since the choice of database can impact runtime performance, MetaPro’s ability to use customized databases offers the potential for the user to select smaller and more specialized databases that would result in significant performance speedups. While MetaPro’s default use of large databases, and multiple passthroughs of data using different tools slows performance, it does ensure both enhanced coverage and improved accuracy of taxonomic and functional (i.e. EC annotations) assignments.

### Easy installation of MetaPro through Docker

Data analysis pipelines often require many 3^rd^-party tools being used in succession, with each tool requiring installation before the pipeline can be deployed. The challenge associated with installing and maintaining tools can represent a barrier to adoption. To overcome these concerns, MetaPro is embedded within a Docker ‘image’^66^, ensuring that the pipeline is deployed with minimal intervention required by the user. Docker was selected over traditional environmental deployment methods, such as a package installation, or other virtualization tools such as Virtualbox^67^, because the process is less invasive to the user and renders the pipeline easier to distribute. Docker and Virtualbox behave similarly, in that each create a separate environment in a computing system for specialized work. However, the intended uses are different. Virtualbox is intended for creating an operating system (OS) within an existing OS. This method effectively isolates the Virtualbox from the rest of the system. Docker applies an overlay to the existing OS. This difference is important, as MetaPro was designed to work on large computing clusters administrated externally. In a computing cluster, a specialized Docker implementation called Singularity^28^ is used. Singularity is purpose-built for scientific computing and is deployed as the solution for our specific compute cluster. Virtualbox’s methods make it undesirable on shared computing resources as Virtualbox requires full control of the temporary resource, taking away control from the administrator, and assigning it instead to the user. With Docker/Singularity, the setup is relatively simple; the user only needs to download the docker image and start the environment. Docker images are distributed and maintained through the Docker repository.

### MetaPro online tutorial

To introduce and guide researchers through the process of analysing metatranscriptomic data, MetaPro features a dedicated tutorial mode. The tutorial has been designed for both computational novices as well as more seasoned bioinformaticians starting out in metatranscriptomic analysis. The interactive tutorial takes users through the tasks of filtering data, aligning reads to databases, and scanning identified genes through taxonomic and enzyme classification tools. The tutorial includes additional intermediate steps, such as reading the quality of the data, and performing sanity checks on the data, to highlight the importance of each step and help the user understand how MetaPro parses the data. The tutorial is run within the MetaPro docker container and was developed over several years of workshops and classes provided to both undergraduate and graduate students, as well as through the Canadian bioinformatics workshop series (https://www.bioinfromatics.ca).

## CONCLUSIONS

Increasingly, microbiome studies are shifting emphasis from identifying the composition of complex microbial communities to understanding how they function. Supporting this shift is the emergence of whole microbiome RNASeq (metatranscriptomics). However, despite the recognized value of these datasets, few dedicated tools are currently available for their analysis. To address this need we developed MetaPro, a single package capable of processing and analyzing metatranscriptomic datasets. Our benchmarking analyses show that MetaPro delivers superior performance over existing pipelines in terms of gene, taxonomic and enzyme annotations. Output is delivered as normalized gene expression profiles in terms of RPKM, together with a novel visualization framework based on the network visualization tool, Cytoscape, that allows the display of enzyme expression and the taxa responsible in the context of individual metabolic pathways.

MetaPro is readily installable with limited dependencies allowing deployment across multiple compute architectures, including cloud computing environments, allowing for massively parallel scale up required for the hundreds of millions of reads typically generated in a single experiment. MetaPro is designed to be flexible to account for the incorporation of improved algorithms as they are developed. To help users understand the various steps involved in metatranscriptomic analysis, we provide an established tutorial that has been developed with non-specialists in mind. Currently, the only major limitation to metatranscriptomic analysis using the MetaPro pipeline is speed of execution relative to existing tools. However, we consider speed to be secondary to accuracy. We are currently developing novel methods to enhance pipeline performance in terms of speed without compromising accuracy of the results.

## METHODS

### MetaPro Workflow

The MetaPro workflow is involves a series of steps which first filters, then annotates the sequence data. In initial preprocessing steps, MetaPro identifies and removes low-quality reads and adapters, as well as reads belonging to the host and other potential contaminants. Next reads associated with non-coding RNAmoieties (e.g. rRNAs and tRNAs) are filtered. The remaining, putative mRNA reads are subsequently into contigs, prior to annotation. After this pre- processing, MetaPro begins the annotation process by identifying which genes the reads belong to, using sequence aligners. Once complete, the pipeline attempts to assign the taxonomy of each read while also attempting to annotate enzymatic functions to each gene. Finally, MetaPro summarizes the details of its analysis into a series of output files (**Figure 1**).

### Sequence Preprocessing

MetaPro first identifies and removes segments of reads associated with adapters and low-quality bases at the 3’ end using AdapterRemoval v2.1.7^68^. The number of threads used is set to the maximum allowable number of cores the computing environment provides. This is automatically detected by MetaPro. The trimqualities flag is used, which trims the 5’/ 3’ termini of reads with the quality scores of up to the AdapterRemoval default quality minimum of 2. Other settings rely on default values. If the data is paired-end, the reads are merged using VSEARCH v2.7.1^69^ with the flag -fastq_mergepairs. The data is then processed using a custom script, taking the reads that still have a forward and reverse pairing that meet the quality thresholds. Reads that meet the quality thresholds but missing their paired counterpart are grouped into a collection of ‘singletons’. Low-quality reads that do not meet the quality thresholds are removed using the -fastq_filter of VSEARCH v2.7.1 with the default error threshold score. Next MetaPro provides the user with an option for filtering contaminant sequences and sequences derived from any host organism that might be associated with the sample (for example if the sample was collected from a mouse or human intestinal sample). This is defined through a configuration option set by the user before launching the pipeline.

To identify sequence artifacts arising from library preparation, MetaPro utilizes the UniVec_Core dataset comprising known sequencing vectors, sequencing adapters, linkers, and PCR Primers derived from the NCBI UniVec_Core Database^35^. Host organism sequences to be filtered are provided as a single FASTA formatted file by the user and identified through sequence similarity searches using BWA 0.7.17^32^ and pBLAT 2.0^33^. Due to pBLAT’s inability to support paired-end data, MetaPro further sorts the reads with internal logic. If either one of a pair of reads matches a sequence in the contaminants file, both reads are filtered from downstream analysis. This behaviour can be toggled to a more permissive rule where pairs are retained if either one of the pair does not match a sequence in the contaminants file.

Having identified and removed low quality and contaminating reads, MetaPro next filters for reads of rRNA origin. In a typical RNA extract, 90% or more of reads can be of rRNA origin^70^. While rRNA depletion kits can significantly reduce the proportion of such reads, metatranscriptomic pipelines need to filter for any remaining reads of rRNA origin. rRNA filtering represents one of the most computationally expensive stages of the MetaPro pipeline. To accelerate this step, MetaPro first removes duplicate reads prior to filtering and repopulates the data after filtering. To further increase efficiency, rRNA filtering is first performed by Barrnap, a rRNA filtering tool based on nhmmer^71^, and then by Infernal^48^, which while more computationally intensive, provides greater sensitivity through the application of covariance models based both on sequence and secondary structure comparisons. To better exploit parallel computing environments, MetaPro splits read data into smaller chunks consisting of 50,000 reads. Each chunk is then processed by Barrnap in parallel, after which the pipeline segregates reads into putative mRNA reads and other reads. The putative mRNA reads are then processed a second time by Infernal and reads not identified as mRNA are removed.

To improve speed and accuracy of the subsequent annotation steps, the pipeline attempts to assemble reads into longer contiguous sequences (‘contigs’) using the sequence assembler, rnaSPAdes^30^. Assembling reads into contigs reduces the amount of file reading and writing (I/O) related tasks by shrinking the number of unique segments analyzed. Furthermore, by analyzing longer sequences, the subsequent alignments of the remaining data produce results of higher confidence. As contigs may represent transcriptional units containing multiple genes, MetaPro applies MetaGeneMark v1^31^, a gene model predictor, to separate contigs into individual putative genes.

### Gene Annotation

Following the data preprocessing stages, the MetaPro pipeline attempts to annotate the putative mRNA transcripts (putative genes derived from MetaGeneMark and unassembled singletons) to known genes. To achieve this MetaPro uses a three-step process to annotate the putative mRNA transcripts to genes. The first step uses BWA 0.7.17^32^ to identify sequence similarity matches to genes in the ChocoPhlAn database^26^. We consider a high confidence annotation at this step to be a match with over 90% sequence identity over 90% of the read length. Next, pBLAT 2.0^33^ is used on unmatched transcripts to identify less stringent sequence similarity matches to genes in the ChocoPhlAn database using a cut-off of >85% sequence identity over at least 65% of the length of the putative mRNA transcript, with a match score of >60. Finally, any unmatched putative mRNA transcripts are processed by DIAMOND 0.9.19^34^ to identify significant sequence similarity matches to sequences collated by the NCBI non-redundant protein database (NR). As for pBLAT, we apply a cut-off of >85% sequence identity over at least 65% of the length of the putative mRNA transcript, with a match score of >60. In situations where a putative mRNA transcript is annotated to multiple genes or proteins, the match-quality score is used to resolve the ambiguity. For transcripts aligned by BWA, the match quality score is the alignment score, under the heading “AS”. In pBLAT and DIAMOND, the match quality score is the bitscore. If a transcript is aligned to multiple different genes or proteins, the highest score among the alignments is selected. In cases where there is a tie in the match quality score, the first result is taken for annotation purposes. The results from each step of this annotation process are compiled into a human readable mapping of genes to the sequences in the dataset annotated to them for use in subsequent stages. Genes identified from the searches of the ChocoPhlAn database are translated into proteins for downstream enzyme annotation.

As for the rRNA filtering step, to make better use of parallel computing, MetaPro splits the data into smaller chunks comprising 50,000 sequences. Only when all BWA searches have been performed will the process move to pBLAT and likewise, only when all pBLAT searches have been performed will the process more to DIAMOND.

### Enzyme Annotation

After annotating putative mRNA transcripts to genes, then converting those genes to proteins, MetaPro will attempt to annotate both the translated genes and the proteins identified by DIAMOND searches of NR, to enzymatic functions, as defined by Enzyme Classification (EC) identifiers. This involves a combination of the enzyme profile tools, DETECT^36^ and PRIAM^37^, together with DIAMOND searches against the Swiss-Prot database^72^. The results are combined by prioritizing the annotation from DETECT and, in cases where enzymatic function could not be assigned using DETECT, enzymatic functions that are consistently assigned by both DIAMOND and PRIAM. This process may assign multiple ECs per transcript. Proteins with multiple EC assignments are further filtered to include only ECs that have been observed to co-occur in the Swiss-Prot database. MetaPro exports two EC reports; low-quality and high-quality. Both EC reports will contain all results from DETECT together with the intersection of predictions from PRIAM and DIAMOND, with an e-value cut-off of <1e-5 for the low-quality report, and an e-value cut-off of <1e-10, together with a PRIAM probability score of >=0.5 for the high quality report.

### Taxonomic Annotation

The MetaPro pipeline uses a consensus of three strategies to assign reads to taxonomic classifications. Firstly, the taxonomic identification numbers (‘taxid’) for each gene/protein for which putative mRNA reads were previously annotated are retrieved based on accession numbers derived from the NCBI accession2taxid database^35^. Secondly, all putative mRNA sequences are processed by the short-read classifiers Kaiju^39^ and Centrifuge^40^. Taxonomic classifications from each source (NCBI lookup, Kaiju and Centrifuge) are then merged, using the classification consensus tool, WEVOTE^41^. Since we expect NCBI lookup-based annotations to be of higher quality, MetaPro assigns greater weight to the NCBI lookup assignments (60% NCBI lookup, 20% Centrifuge, and 20% Kaiju). This effectively results in the use of Kaiju and Centrifuge to cover potential gaps in annotation.

### Output and Visualizations

#### Read metrics and data quality

MetaPro compiles various read metrics within a text file. This file reports: the total number of sequence reads from the raw input, the remaining number of high-quality reads after filtering for quality and adaptors, the number of reads associated with the host (if applicable), the amount of rRNA and tRNA reads removed from the rRNA filtration step, the number of putative mRNA reads, the number of reads annotated to a gene or protein, the total number of unique transcripts found, and the number of unique enzymes detected in the data. MetaPro also creates a histogram of the number reads and their quality scores, before and after they have been filtered for low-quality reads. Finally, the pipeline reports on the N50 and L50 read statistics of the contigs formed during the assembly stage.

#### Gene expression

MetaPro produces a collection of genes, along with their constituent reads, summarized in a gene/protein-to-read map. This gene/protein map labels each gene (ChocoPhlAn) or protein (NR), along with all the read IDs of the input data that annotate to that gene/protein. MetaPro additionally provides a table of genes/proteins, associated EC, and RPKM, as well as another table of genes/proteins, with their full taxonomy. MetaPro also produces a contig-to-read map, that shows the reads associated with each contig generated by rnaSPAdes.

#### Enzyme annotation

To add context to the enzyme annotation results, MetaPro exports a table file that is compatible with Cytoscape^73^ import functions. The table provides a list of ECs, and the expression (as measured by RPKM) of the various genes and proteins that identify to that EC. EC expression is further broken down by taxa, selected on the basis of minimal abundance. Specifically for each taxon, starting at the rank of species, if that taxon is not associated with at least 1% of total reads (default cut-off) then it is merged with other taxa that share the genus that also do not exceed the 1% criteria. This process is repeated to define groups of taxa at the level of family, order, class and phyla, that represent at least 1% of total reads. Together with the installation of two Cytoscape apps (KEGGscape and enhanceGraphics), KEGG defined metabolic pathways can be imported as KGML files and enzymes in the pathways annotated with the MetaPro defined EC file. In typical applications, this can result in the representation of each enzyme as a pie chart or annular ring, in which the size of the pie chart/ring represents total expression (as defined by RPKM values) of that EC in the dataset and the segments indicate the taxa responsible for contributing to the expression of that enzyme. In addition, MetaPro also generates a table of ECs, grouped together by super-pathways (EC_coverage.csv) and a table of super-pathways, and their constituent ECs, expressed as RPKM for the formation of the EC super-pathway heatmap image.

#### Taxonomic annotation

MetaPro reports the taxonomic findings of the sample as a table of taxa and read counts. In brief, annotated genes and proteins are assigned to and grouped on the basis of taxonomic assignment. Reads assigned to each gene/protein that belongs to a specific taxon are summed to provide to produce a read count for each taxa.

### Computing considerations

Since cluster computing environments can impose limitations on processing time available to the user, MetaPro was developed to improve data throughput through parallelization and ensure the completion of the pipeline through the implementation of an auto-resume feature. To address the former, for the rRNA filtration and gene annotation steps, we deploy a technique based on MapReduce^74^ in which datasets are divided into smaller chunks of 50,000 sequences that are processed on parallel threads. Other steps were found not to benefit from splitting the dataset and are simply run as serial processes on the global dataset. Since the processing time required by MetaPro is typically unknown before runtime, to ensure that the processing of a dataset is completed, particularly in computing environments where allocations may be time limited, MetaPro features a mutli-level auto-resume feature. This is enabled through a robust bookmarking system that keeps track of the state of processing prior to termination due to time constraints. Subsequent restarts allow MetaPro to resume processing on the remaining sharded data (i.e. for the rRNA filtering and gene annotation steps). For other steps, MetaPro will restart from the beginning of the stage that failed.

### Initial Set-up

MetaPro features configurable parameters for many features that impact processing runtime. In a master configuration file, users define the locations of the library and reference file paths required by the various tools employed by the pipeline. The configuration file also includes a series of settings to define stringency levels of filters and displays such as: the cut-off for defining taxa to be reported in the enzyme annotation output file; the CIGAR match length cutoff for BWA gene annotation; and the identity, length, and score cutoffs for accepting pBLAT and DIAMOND gene annotations. The configuration file lets users control the efficiency of the parallelization, by adjusting the size of the data chunks processing in the rRNA filtration and gene annotation steps. The user can also control the number of concurrent processes running at each step, and the memory allocated for rRNA filtering, and gene annotation. Finally, the configuration file lets users control the state of the interim files; interim files can be retained intact, compressed, or removed to save disk space.

## ACKNOWLEDGEMENTS

We would like to thank Ana Popovic, Emil Jurga, James St Pierre and Donny Chan for their enthusiastic help in pipeline testing and providing valuable feedback on pipeline implementation. We would also like to thank Ben Jackel for advice on analysing output generated by HUMAnN2 and HUMAnN3.

## CONTRIBUTIONS

BT, MA and XX developed the pipeline. NN, JA and JP helped in the design of the pipeline. BT, MA and JP wrote the manuscript. JP conceived the project.

## CODE AVAILABILITY

The software is freely available under the GNU public license V3 and can be accessed through https://github.com/ParkinsonLab/MetaPro.

## SUPPLEMENTAL FIGURES

Supplemental Figure 1: Gene annotation performance of MetaPro, HUMAnN3, and HUMAnN2

Stacked barcharts depicting the number of reads annotated to specific taxa in (A) NOD mouse samples, and (B) Kimchi samples by BWA alignments, MetaPro, HUMAnN3, and HUMAnN2. The NOD mouse datasets were generated from gut samples from mice inoculated with a defined microbial consortium (Altered Schaedler Flora (ASF); ^42^). In addition to the 8 taxa associated with ASF, reads were also assigned to *Parabacteroides goldsteinii,* a close relative of *Parabacteroides ASF519* (see legend). The kimchi datasets comprise five major taxa (see legend; ^75-79^). It should be noted that *Leuconostoc gasicomitatum* reported in the original publication is currently classified as a subspecies of *Leuconostoc gelidum*. For NOD sample SRR1828965, HUMAnN3 did not annotate any reads; for kimchi sample SRR443366, HUMAnN2 did not annotate any reads.

Supplemental Figure 2: Taxonomic classification performance of MetaPro, HUMAnN3, and HUMAnN2

For the NOD mouse (A) and Kimchi (B) datasets, each pie chart shows a breakdown of taxonomic assignments at the different phylogenetic levels indicated, that are closest to the last common ancestor of the expected bacteria within the sample. For the human oral datasets (C), given the lack of gold standard assignments, each pie chart represents the relative abundance of reads assigned to different phylogenetic levels. Unclassified reads represent annotated reads with no assigned taxon. The graphs indicate a substantial improvement in HUMAnN3’s annotation abilities over HUMAnN2.

Supplemental Figure 3: Enzyme annotation performance of MetaPro and HUMAnN2

Stacked barcharts indicate the number of enzymes, as defined through enzyme classification (EC) assignments, annotated by each pipeline for the three sets of datasets: (A) NOD Mouse, (B) Kimchi, and (C) Human Oral Biofilm data. In addition to displaying ECs unique or shared between MetaPro and HUMAnN2, also shown are ECs, predicted by HUMAnN2 to occur in combination with another EC, in the same transcript, with no supporting evidence that such a combination has been previously observed (as defined through Swiss-Prot annotations). Further, for HUMAnN2, we show the number of EC assignments that occur in combinations of three or more ECs.

## SUPPLEMENTAL TABLES

Supplemental Table 1 Summary of Sequence Read Processing for Three Metatranscriptomic Datasets (NOD Mouse gut; Kimchi and Human Oral Biofilm) Processed by HUMAnN3, HUMAnN2, MetaPro and SAMSA2.

This table reports the processing results from the four pipelines on samples from three different datasets. HUMAnN2 and HUMAnN3’s preprocessing tool concatenates paired reads into 1 single file and treats them as 2 separate reads. The NOD mouse samples are paired-ended data, while the kimchi and human oral biofilm represent single-end sequence datasets. Unlike MetaPro and SAMSA2, HUMAnN3 and HUMAnN2 do not report transcripts but instead group proteins identified in their pipelines into gene families that are reported in the final column.

Supplemental Table 2: Read annotation statistics for NOD mouse datasets from MetaPro, HUMAnN3, HUMAnN2, SAMSA2 compared with the gold standard.

This table shows the amount of reads in each NOD mouse sample each pipeline assigned to the 8 ASF bacteria: *Clostridium ASF356, Clostridium ASF502, Eubacterium plexicaudatum, Firimicutes ASF500, Lactobacillus ASF360, Lactobacillus murinus, Mucisprillim schaedleri, and Parabacteroides ASF519*. Due to the similarity *between P. ASF519*, and *P. goldsteinii*, the pipelines will sometimes annotate to *P. goldsteinii* rather than to *P.ASF519*. *P. goldsteinii* was also a dominant species found within the samples outside of the 8 ASF bacteria. The expected results were produced by annotating the reads with a reference containing only the 8 ASF, using BWA.

Supplemental Table 3: Read annotation statistics for kimchi fermentation datasets from MetaPro, HUMAnN3, HUMAnN2, SAMSA2, compared with the gold standard.

This table shows the number of reads in each kimchi sample, annotated to the expected 5 lactic acid bacteria (LAB) from each pipeline: *Leuconostic mesenteroides*, *Lactobacillus sakei*, *Weissella koreensis*, *Leuconostoc carnosum*, and *Leuconostoc gelidum*. The expected results were obtained by annotating the kimchi datasets against a database containing only the reference gene sequences for the 5 LAB, on BWA.

Supplemental Table 4: Comparisons of Enzyme annotations between MetaPro and HUMAnN3 for NOD mouse, kimchi, and human oral biofilm datasets.

This table compares the ECs of MetaPro against HUMAnN3 on the NOD mouse, kimchi, and human oral biofilm datasets. The HUMAnN3 ECs were filtered for EC co-occurrence pairs that were not found in Swiss-Prot, and multiple unique ECs that annotated to the same gene family. The resulting HUMAnN3 ECs were contrasted against MetaPro’s EC, yielding a common set of ECs found in both tools, ECs found only by MetaPro, and ECs found only by HUMAnN3. The same comparison is shown for MetaPro’s high-qaulity EC predictions.

Supplemental Table 5: Comparisons of Enzyme annotations between MetaPro and SAMSA2 for NOD mouse, kimchi, and human oral biofilm datasets.

This table compares the ECs of MetaPro against SAMSA2 on the NOD mouse, kimchi, and human oral biofilm datasets. The SAMSA2 ECs were filtered for EC co-occurrence pairs that were not found in Swiss-Prot, and multiple unique ECs that annotated to the same gene family.

The resulting SAMSA2 ECs were contrasted against MetaPro’s EC, yielding a common set of ECs found in both tools, ECs found only by MetaPro, and ECs found only by SAMSA2. The same comparison is shown for MetaPro’s high-qaulity EC predictions

Supplemental Table 6: Comparisons of Enzyme annotations between MetaPro and HUMAnN2 for NOD mouse, kimchi, and human oral biofilm datasets.

This table compares the ECs of MetaPro against HUMAnN2 on the NOD mouse, kimchi, and human oral biofilm datasets. The HUMAnN2 ECs were filtered for EC co-occurrence pairs that were not found in Swiss-Prot, and multiple unique ECs that annotated to the same gene family. The resulting HUMAnN2 ECs were contrasted against MetaPro’s EC, yielding a common set of ECs found in both tools, ECs found only by MetaPro, and ECs found only by HUMAnN2. The same comparison is shown for MetaPro’s high-qaulity EC predictions.

Supplemental Table 7: Computational performance statistics of MetaPro, HUMAnN3, and SAMSA2

This table reports the amount of processing time required for each run of the three pipelines. MetaPro additionally exports the timing data of each stage independently. HUMAnN3’s pre- processing step is a separate stage using a separate tool called KneadData. SAMSA2 cleans the data in the pipeline, but it is integrated and does not export timing data.

